# A ribozyme mass extinction at the RNA-cellular boundary and its potential imprint on the genetic code

**DOI:** 10.64898/2026.03.05.709948

**Authors:** Ido Bachelet

## Abstract

The transition from the RNA world to cellular life remains one of the least understood events in the history of biology. This study proposes that geochemical changes approximately 3.9-3.8 billion years ago triggered a mass extinction of ribozymes - possibly the first extinction event on Earth - and that the genetic code bears the fingerprints of its survivors. The argument draws on parallels with animal mass extinctions, where generalist species survive, dominate post-extinction ecosystems as disaster taxa, and shape subsequent evolutionary trajectories. Among small self-cleaving ribozymes, the hammerhead ribozyme accounts for approximately 91% of all ∼221,000 known sequences and is the only ribozyme that is found across all kingdoms of life. The phylogeny of hammerhead ribozymes exhibits a star-like topology consistent with rapid post-bottleneck expansion, and its small size, ability to tolerate diverse chemical conditions, and broad substrate specificity suggest it was a resilient generalist feeder fitting as a disaster taxon. Other families of ribozymes possibly survived in small, taxonomically isolated populations resembling relict species in ecological refugia. It is suggested that the body plans of surviving ribozymes seeded the primitive processes that would later become the genetic code, for example with RNA-degrading trinucleotides becoming stop codons, partitioning the trinucleotide space into signals of termination and translation. This hypothesis proposes a reframing of the origin of the genetic code in part as an ecological legacy rather than a purely chemical inevitability.

## Introduction

The RNA world hypothesis posits a period in Earth’s early history during which RNA molecules served simultaneously as genetic material and catalytic agents ^1,2^. This hypothesis, grounded in the discovery of catalytic RNA ^3,4^, is now broadly accepted as a foundation of origin-of-life theory. Yet the transition from the RNA world to the cellular world remains one of the least understood events in the history of life ^5,6^.

Most accounts of this transition emphasize its constructive aspects: the synthesis of fatty acid vesicles, the encapsulation of replicating molecules, and the emergence of protocells with rudimentary metabolism ^7-9^. Less discussed is the possibility that this transition was also destructive - that the geochemical changes enabling cellular life simultaneously devastated the pre-existing RNA ecosystem. In the animal kingdom, the most consequential evolutionary transitions are routinely associated with mass extinction events ^10,11^. The end-Permian extinction cleared the way for dinosaur radiation ^12^; the end-Cretaceous extinction enabled the rise of mammals ^12,13^. In each case, what appears as a creative explosion was preceded by catastrophic destruction.

This paper proposes that the RNA-to-cellular transition was triggered by a mass extinction of ribozymes - potentially the first mass extinction event on Earth - and that the genetic code bears the chemical fingerprints of its dominant survivor. The argument proceeds in three steps. First, geochemical evidence is reviewed for environmental changes ∼3.9-3.8 Ga that would have been catastrophic for free-living RNA organisms. Second, evidence is presented that extant ribozyme families exhibit every hallmark of a post-mass-extinction ecosystem, with the hammerhead ribozyme as the disaster taxon. Third, computational analysis demonstrates that the trinucleotide composition of the ribozyme body plan regions maps onto the functional architecture of the genetic code.

## Results

To identify candidate triggers for a possible major ecological shift occurring at the RNA-cellular (R-C) boundary, we surveyed the geochemical literature for changes proposed to have taken place between approximately 4.2 and 3.5 Ga. Four interlinked factors converge within the interval ∼3.9-3.8 Ga **(Fig. 1)**. The upper bound of this interval is constrained by geochemical modeling showing the onset of rapid environmental change, and the lower bound by the earliest isotopic signatures of biological carbon fixation at ∼3.8 Ga ^14,15^, indicating that cellular life was already emerging by that time. First, the late accretion bombardment declined sharply across this interval ^16^ **(Fig. 1A)**. While the waning of impact sterilization is typically viewed as permissive for life, it simultaneously reduced the meteoritic delivery of metals and reduced phosphorus species that had sustained prebiotic chemistry ^17,18^. Second, surface temperatures plunged from over 200°C to near-modern values across this interval **(Fig. 1B)**. While the cooling itself would have expanded the thermal range habitable by RNA, it accelerated silicate weathering of the evolving crust, releasing cations into the ocean and drawing down atmospheric CO_2_ - directly driving a rapid pH rise ^19,20^. Ocean pH rose from approximately 5.0 toward near-neutral values more rapidly than previously estimated ^21^ **(Fig. 1C)**. The sensitivity of RNA to alkaline hydrolysis is intrinsic to its chemistry ^22^, hence rising pH would have progressively shortened the half-life of exposed ribozymes. Fourth, the delivery of reduced phosphorus by late accretion declined by orders of magnitude across this interval ^17,23^ **(Fig. 1D)**, while phosphate sequestration through mineral precipitation further constrained its availability ^18,24^. Ribozyme populations thus faced a growing scarcity of their fundamental building blocks. These four stressors are not independent: the declining bombardment drove the thermal crash, which in turn accelerated pH equilibration through enhanced weathering, which promoted phosphate mineralization. The result was a synergistic collapse - ribozymes simultaneously lost structural stability, catalytic competence, and RNA building blocks (food) for reproduction. This convergence of stressors mirrors the multi-causal structure of animal mass extinctions, in which synergistic interactions between environmental changes amplify extinction severity beyond what any single trigger could produce ^25,26^.

**Figure 1.**
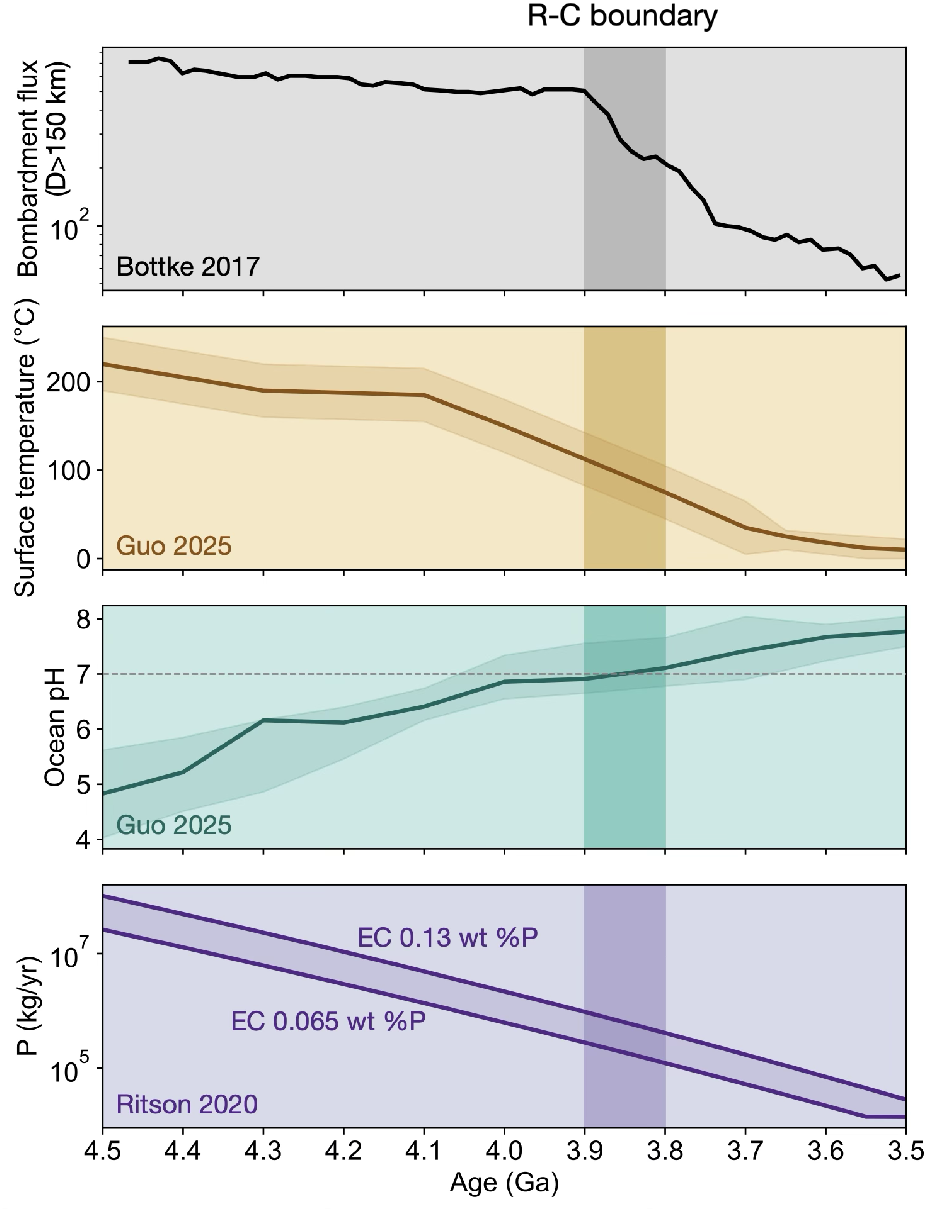
Geochemical changes at the RNA-cellular boundary. Composite timeline of environmental variables across the 4.5-3.5 Ga interval, with the proposed R-C boundary interval (∼3.9-3.8 Ga) indicated. (A) Declining impact flux (data from Bottke & Norman, 2017). (B) Surface temperature evolution showing the crash from >200 °C to near-modern values (data from Guo & Korenaga, 2025). (C) Ocean pH rise from acidic to neutral (data from Guo & Korenaga, 2025). (D) Declining delivery of reduced phosphorus by late accretion (data from Ritson et al., 2020). Together, these panels document a convergence of environmental stressors - physical (bombardment), thermal (cooling), chemical (pH shift), and nutritional (phosphate depletion) - coinciding with the proposed extinction interval.

A key detail in understanding the R-C transition is the distribution of extant ribozyme families, which exhibits a pattern strikingly consistent with a post-mass-extinction ecosystem **(Fig. 2)**, with each hallmark finding a direct parallel in the animal mass extinction record **(Table 1)**. Of approximately 221,000 known self-cleaving ribozyme sequences across all ten families, approximately 199,000 (91%) are hammerhead ribozymes ^27,28^ **(Fig. 2A)**, a degree of numerical dominance characteristic of disaster taxa ^29^, exceeding even the post-Permian dominance of *Lystrosaurus* (70-95% of terrestrial vertebrate individuals) ^30,31^. The other types of small, self-cleaving ribozymes exist in taxonomically isolated populations (82 to ∼11,500 sequences) consistent with relict species in ecological refugia ^32,33^. The hatchet ribozyme (∼166 copies restricted entirely to *Crassvirales* phage genomes) represents an extreme case of refugial survival, paralleling the coelacanth ^34^. The hammerhead satisfies many criteria of a disaster taxon: it is a small, adaptive (modular) ^35^ and versatile ribozyme; it can tolerate and operate in diverse, non-standard chemical conditions ^36,37^; and it possesses the broadest substrate specificity (NUH↓, covering 12 of 16 dinucleotide combinations) of all small, self-cleaving ribozymes.

**Table 1.** Parallels between ribozyme and animal mass extinction signatures. Each row identifies a hallmark signature of mass extinction and recovery, with the ribozyme evidence from this study and its closest paleontological analog. Ribozyme citations refer to independent published studies except where noted; paleontological analogs span the end-Permian, end-Cretaceous, and Late Ordovician mass extinctions. See Supplementary Note S2 for expanded discussion of each parallel.

| **Extinction signature** | **Ribozyme extinction** | **Paleontological analog** |
| --- | --- | --- |
| Disaster taxon dominance | Hammerhead: ~91% of ~221,000 sequences; present in all domains of life<br>De la Peña & García-Robles 2010; Perreault et al. 2011 | *Lystrosaurus*: 70-95% of Early Triassic terrestrial vertebrate individuals<br>*Botha-Brink et al. 2016; Kauffman & Harries 1996* |
| Relict in refugium | Hatchet: ~166 copies restricted to Crassvirales phage genomes<br>Rfam RF02678; Weinberg et al. 2019 | *Latimeria (coelacanth)*: 2 species in deep-sea caves; presumed extinct 66 Myr<br>*Smith 1939; Amemiya et al. 2013* |
| Relict survivor (restricted range) | Hairpin (~139), Pistol (~103), Twister sister (~129): each in narrow host ranges<br>Rfam 15.1; Weinberg et al. 2019 | *Nautilus*: sole extant nautiloid; restricted Indo-Pacific after ammonite extinction<br>*Ward 1987* |
| Generalist survival advantage | Hammerhead NUH↓: 12/16 dinucleotide targets (NHH rule); tolerates clay, Fe<sup>2+</sup>, extreme conditions<br>Kore et al. 1998; Athavale et al. 2012; Biondi et al. 2007 | K-Pg bivalves: genera in ≥3 biogeographic provinces had ~20% extinction vs. 70% for restricted<br>*Jablonski 2005* |
| Marine disaster taxa | Hammerhead dominant across marine invertebrate host phyla (Trematoda, Bivalvia, Cephalopoda)<br>De la Peña & García-Robles 2010; Perreault et al. 2011 | *Claraia, Unionites, Lingularia*: low-diversity high-dominance Early Triassic benthos<br>*Clapham et al. 2016* |
| Star phylogeny / post-bottleneck radiation | Hammerhead: star-like tree across 97 genera; D = +3.79, Fs = -690.8<br>This study | Neoaves: explosive diversification at K-Pg; unresolved star polytomy in molecular phylogenies<br>*Jarvis et al. 2014* |

**Figure 2.**
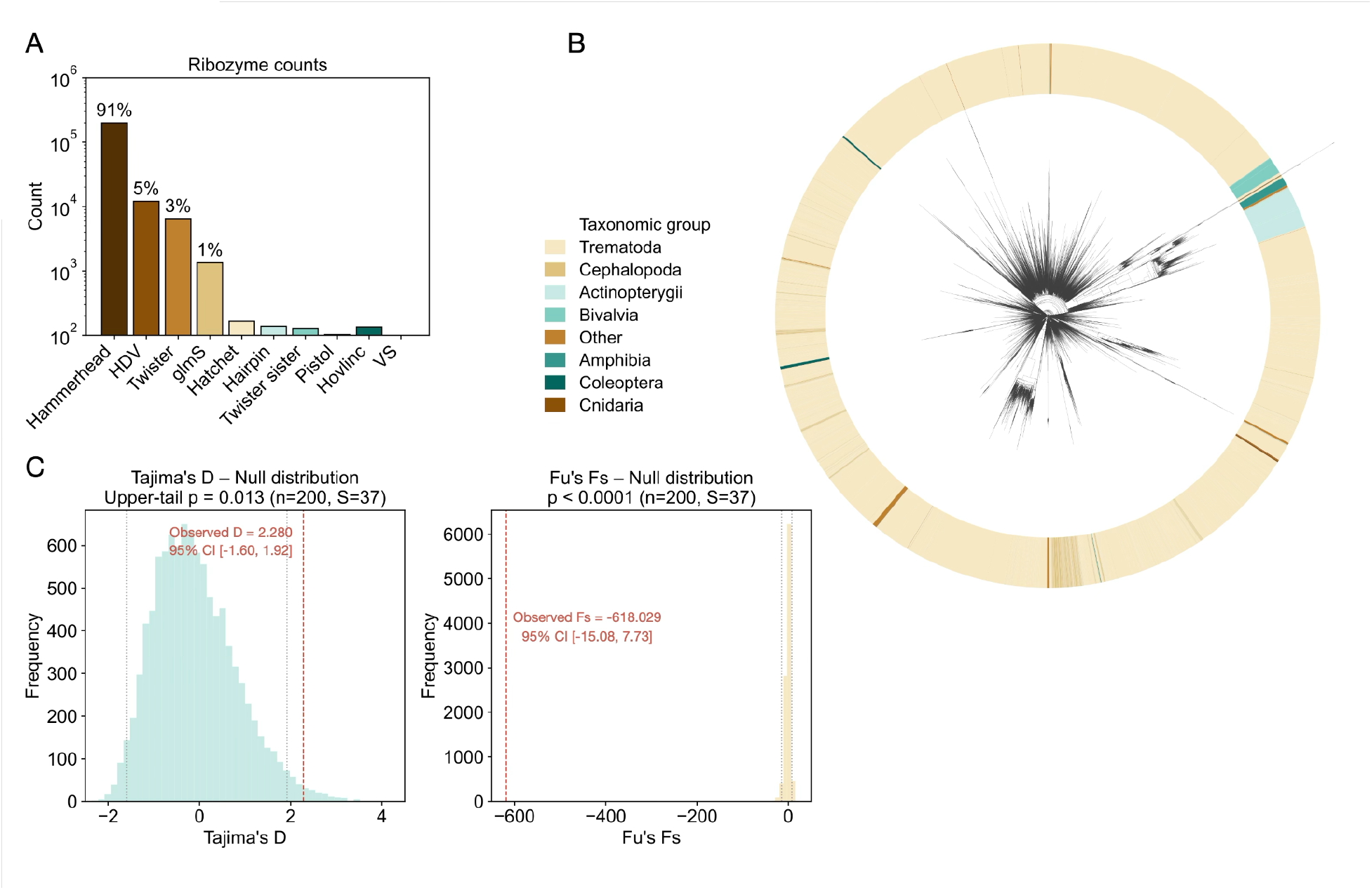
Evidence for ribozyme mass extinction. (A) Abundance distribution across all ten known small self-cleaving ribozyme families, showing the hammerhead’s 91% dominance, a pattern characteristic of disaster taxa following mass extinction events. (B) Circular phylogeny of 80,548 unique hammerhead sequences colored by host taxonomic group. The star-like topology is consistent with post-bottleneck radiation, and the interspersion of distantly related host phyla around the tree reflects repeated independent colonization events rather than co-speciation, consistent with horizontal transfer of a single dominant survivor lineage. (C) Null distribution tests for Tajima’s D (left) and Fu’s Fs (right), each based on 10,000 coalescent replicates (n = 200, S = 40). Blue and orange histograms show the null distributions under neutral expectations; red dashed lines indicate the observed values (D = +2.28, upper-tail p = 0.013; Fs = −618.0, p < 0.0001). Observed D falls in the extreme upper tail (98.7% of null replicates below observed), indicating deep hierarchical structure; observed Fs falls far below any null replicate, indicating massive haplotype excess. The discordance between positive D and extremely negative Fs is diagnostic of ancient divergence preserved through a bottleneck followed by rapid within-lineage expansion. Full-sample values (n = 5,036) are D = +3.79, Fs = −690.8.

Disaster taxa expand in a typical pattern. In order to examine whether hammerhead ribozymes exhibit such a pattern, the phylogeny of 80,548 unique complete hammerhead sequences spanning 97 genera was constructed. This phylogeny exhibits a star-like topology consistent with rapid post-bottleneck expansion **(Fig. 2B)**. Population genetics statistics computed on the full stratified subsample (n = 5,036; S = 40) reveal Tajima’s D = +3.79 and Fu’s Fs = −690.8; coalescent null testing on a matched subsample (n = 200; S = 37; 10,000 replicates) confirms both values are extreme (D = +2.28, upper-tail p = 0.013; Fs = −618.0, p < 0.0001) **(Fig. 2C)**. The discordance between positive D (reflecting deep hierarchical structure across 97 genera) and extremely negative Fs (reflecting haplotype excess from rapid within-lineage expansion) is diagnostic of ancient divergence preserved through a bottleneck, followed by explosive radiation within surviving host lineages ^38^.

The mass extinction of ribozymes placed the hammerheads that survived it at a unique position that allowed these ribozymes to shape the early exit from a pre-code world. The emergence of the genetic code is one of the most profound questions in biology, and is still under active study ^39-42^. It has been addressed by various angles, including chemistry and information theory, but rarely as a legacy of ecological circumstance at the R-C boundary. Some of the predictions of this hypothesis were next tested. If the hammerhead’s body plan shaped part of the emergence of the genetic code, it would have been likely that the functional organs of the hammerhead partitioned the trinucleotide sequence space in ways corresponding to functional categories of the genetic code.

We first analyzed stem and core regions from the entire curated dataset of hammerhead ribozymes to see if such partitions exist. Two results confirmed this hypothesis. Stems and cores contain RNA triplets that clearly segregate by their roles in the genetic code **(Fig. 3A)**. In particular, amino acids that are hypothesized to have entered the genetic code early, are enriched 2.8-fold in stems (0.8-0.9-fold in cores) **(Fig. 3B)**. A Spearman correlation between codon-level stem enrichment and amino acid chronological rank yielded ρ = −0.243 (p = 0.06, permutation p = 0.056): marginal, but in the predicted direction. The correlation is driven by the binary partition between core-resident and non-core trinucleotides rather than a smooth gradient, reflecting the discrete nature of ribozyme architecture.

**Figure 3.**
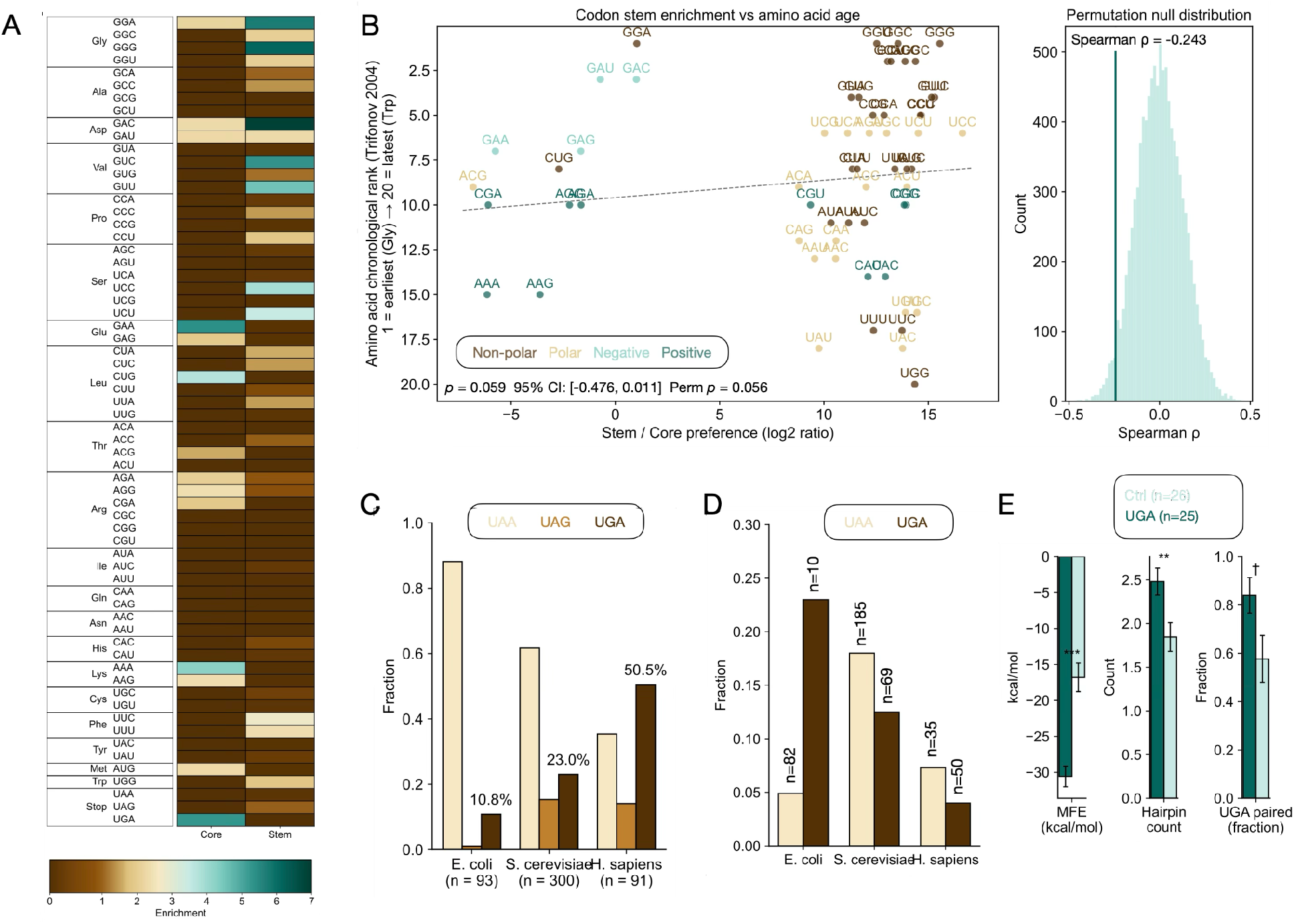
The body plan’s imprint on the genetic code. (A) Trinucleotide enrichment heatmap showing stop codons enriched in catalytic cores (6.1-fold) and early amino acid codons enriched in structural stems (2.8-fold for Val/Asp/Gly). (B) Stem enrichment vs. amino acid chronological rank (Trifonov consensus), showing a marginal but predicted negative correlation at codon level (ρ = −0.243, p = 0.06, permutation p = 0.056). (C) Stop codon usage and downstream context in universally conserved genes across *E. coli, S. cerevisiae*, and *H. sapiens*. Left: fractional usage of UAA, UAG, and UGA, showing a progressive shift from UAA dominance in *E. coli* (88% UAA) to UGA dominance in *H. sapiens* (51% UGA), consistent with UGA’s gradual co-option from a pre-translational cleavage signal into a translational stop codon. (D) Mean information content (IC) of the downstream hexamer for UAA-vs. UGA-terminated universal genes, showing that UAA-terminated genes carry higher downstream sequence constraint in *E. coli* and *S. cerevisiae*, consistent with UAA having been under translational selection longer than UGA. (E) UGA readthrough vs. termination structural analysis. Readthrough sites (n = 25; 20 selenoprotein SECIS elements, 5 cellular readthrough genes) show significantly more stable local RNA structure than UGA termination controls (n = 26; MFE: −30.6 vs. −16.8 kcal/mol, p = 1.2 × 10^−5^) and more stem-loop structures (2.48 vs. 1.85 hairpins, p = 0.006), consistent with local structure shielding UGA from release factors and preserving its accessibility to alternative decoding machinery.

Interestingly, standard stop codons are enriched 3.4-fold in catalytic cores of all ribozymes, and 6.1-fold in the hammerhead core specifically, while being depleted 0.4-fold in stem regions **(Fig. 3A, Fig. S1)**. The rationale for catalytically-active RNA triplets assuming the role of stop signals is twofold: first, they are hazardous to the integrity (and thus reliability) of RNA messages, so they need to be kept as far as possible from the message (at its end); and second, they are by definition natural terminators of RNA strands.

In a wide range of hammerhead ribozymes, the most striking feature of the catalytic core is the motif CUGAUGA, which includes two tandem repeats of the UGA (opal/umber) stop codon. The hammerhead is the only of the small self-cleaving ribozymes to contain two stop codons. The unique placement of the hammerhead in the post-extinction world raises the hypothesis that UGA was the first primitive stop codon, and was replaced only later by UAA. UGA’s leakiness and frequent reassignments ^43,44^ could be reinterpreted not as recent recruitment but as evidence of its pre-translational origin; it was never optimized for the release factor system. This is supported by analysis of 492 universally conserved genes across three organisms: universal genes strongly prefer UAA (e.g., *E. coli*: 88% vs. 64% background, p = 5.4 × 10^−6^), consistent with UAA as the ancestral translational stop. In *S. cerevisiae* and *H. sapiens*, UGA shows less downstream context bias than UAA (*S. cerevisiae*: mean hexamer IC = 0.12 vs. 0.19 bits; *H. sapiens*: 0.06 vs. 0.06 bits), consistent with UGA’s function originating in a pre-translational system where downstream context was irrelevant ^45,46^ **(Fig. 3C, 3D)** Furthermore, UGA readthrough sites - where UGA is decoded as selenocysteine or other amino acids rather than functioning as a stop - show significantly more stable local RNA structure than efficiently terminating UGA controls (MFE: −30.6 vs. −16.8 kcal/mol, p = 1.2 × 10^−5^) and more stem-loop structures (p = 0.006), consistent with structural elements shielding UGA from the termination machinery and preserving its pre-translational accessibility to alternative decoding **(Fig. 3E)**.

## Discussion

The central claim of this paper is that the transition from the RNA world to cellular life was triggered by a catastrophic event leading to mass extinction of ribozymes, and that the survivors of this event had left a legible imprint on the genetic code. The evidence presented - geochemical convergence of stressors, ecological signatures of mass extinction among extant ribozymes, and the correspondence between the ribozyme body plan and genetic code primitives - lends support to this hypothesis.

Several aspects of this framework warrant further discussion. The mass extinction model makes a specific prediction about the nature of the geochemical trigger: it should have been rapid relative to evolutionary timescales, multi-causal rather than attributable to a single stressor, and selectively destructive of ecologically specialized forms. The convergence of reduced bombardment, temperature shift, pH changes, and depletion of available phosphate documented for the 3.9-3.8 Ga interval satisfies all three criteria. While individual parameters remain subject to considerable uncertainty - early ocean pH estimates range from 5 to 7 depending on the model [Guo & Korenaga, 2025; Krissansen-Totton et al., 2018; Zahnle et al., 2010] - the convergence of multiple independent environmental shifts is robust across models.

The distinction between ecological stop (UGA) and translational stop (UAA) proposes a reframing of the traditional view of UGA as a late addition to the code ^47^. The present model pictures UGA as the primary weapon carried by the ribozymes that survived the extinction, who carried it in two copies each; it was in this capacity as a ribozyme-degrading blade, that UGA functioned as a natural termination signal for any primordial translation process that utilized RNA as its message and occurred in its vicinity. This might be a reason why the first amino acids to be translated by RNA were translated at the stem, physically remote from this hazardous triplet. The apparent unreliability of UGA in the modern code reflects its ancestry - a signal from a fundamentally different world, which was never fully adapted to the modern system it was grandfathered into.

A further prediction - that UGA readthrough sites would retain structural signatures of their pre-translational history - was confirmed: readthrough sites show significantly more stable local RNA structure than termination controls (p = 1.2 × 10^−5^) and more stem-loop structures (p = 0.006). This is consistent with a model in which UGA’s transition from ecological stop to translational stop was completed most efficiently at structurally accessible sites, where release factors could recognize UGA without steric interference. At readthrough sites, elaborate local structures (SECIS elements, viral pseudoknots) shield UGA from the termination machinery, preserving its availability for alternative decoding - an echo of the post-extinction world in which UGA first acquired biological meaning.

Finally, the present study raises the possibility of a mass extinction event at the R-C boundary, showing multiple parallels to other extinction events from the geological past. Such catastrophic events are known to not only exterminate many clades, but also open new opportunities and trigger rapid evolution and expansion of new branches. It is intriguing to wonder whether, just as the Cambrian explosion followed the Ediacaran extinction, and the mammalian radiation followed the dinosaur extinction, the emergence of cellular life and the genetic code followed the extinction of ribozymes, of which we might learn from their fossil record within genomes across all kingdoms of life.

## Methods

Trinucleotide enrichment analysis. Trinucleotide frequencies were computed for catalytic core and structural stem regions across all seven small self-cleaving ribozyme families using Rfam seed alignments [Kalvari et al., 2021]. Each sequence was parsed into core and stem regions based on published secondary structure annotations. Enrichment was calculated as observed frequency divided by expected frequency (background trinucleotide composition of the full sequence). A total of 524,824 stem trinucleotides and 563,867 core trinucleotides were analyzed.

Pyrimidine-pyrimidine dinucleotide analysis. For each ribozyme family, the frequency of pyrimidine-pyrimidine dinucleotides (CC, CT, TC, TT/UU) was computed in catalytic core sequences and compared to the expected frequency of 0.25 under random composition.

Trifonov consensus chronology correlation. Amino acid chronological ranks were taken from Trifonov’s consensus chronology [Trifonov, 2004]. For each of the 61 sense codons, stem enrichment was calculated as described above. Spearman rank correlation was computed between codon-level stem enrichment and the chronological rank of the corresponding amino acid. Permutation testing (10,000 permutations) was used to assess significance.

Phylogenetic analysis. 80,548 unique complete hammerhead ribozyme sequences were retrieved from Rfam [Kalvari et al., 2021] and aligned using MAFFT v7 [Katoh & Standley, 2013]. An approximate maximum-likelihood tree was constructed using FastTree 2 [Price et al., 2010] with the GTR+CAT model.

Population genetics statistics. Tajima’s D [Tajima, 1989] and Fu’s Fs [Fu, 1997] were computed on a stratified subsample of 5,036 sequences preserving the taxonomic distribution of the full dataset (97 genera). The subsample contained 40 segregating sites and 4,555 distinct haplotypes. P-values were assessed by coalescent simulation (10,000 replicates).

UGA contextual analysis. Stop codon usage and flanking nucleotide context were analyzed in 492 universally conserved genes (ribosomal proteins, translation factors, RNA polymerase subunits) across *E. coli* (93 genes), *S. cerevisiae* (300 genes), and *H. sapiens* (99 genes), compared against 10,298 total coding sequences. Information content was computed for the six positions downstream of each stop codon as IC = Σ p_i_ log_2_(p_i_/0.25).

UGA readthrough analysis. 25 known UGA readthrough sites (20 human selenoprotein SECIS elements: GPX1-4,6, TXNRD1-3, DIO1-3, SELENOP,W,H,I,K,N,S,T, MSRB1; plus 5 cellular readthrough genes: rabbit beta-globin, TMV, MuLV, Sindbis, VEGFA-Ax) were compared against 26 control UGA terminators from housekeeping genes (8 *E. coli*, 4 *S. cerevisiae*, 14 *H. sapiens*). All sequences were fetched from NCBI. Minimum free energy, base-pairing probability, and hairpin loop count were computed using RNAfold from ViennaRNA 2.7.0 [Lorenz et al., 2011] on 103-nucleotide windows (50 nt upstream + UGA + 50 nt downstream). Statistical comparisons used Mann-Whitney U tests (MFE, pairing probability, hairpin count) and Fisher’s exact test (UGA paired/unpaired classification).

Body plan classification. Ribozyme body plan classification follows the framework established in Bachelet [2026], with structural regions designated as bodies (paired stems), cavities (catalytic cores), and limbs (unpaired substrate-interacting strands).

## Author contributions

I.B. designed experiments, performed experiments, analyzed data, and wrote manuscript.

## Competing interests declaration

The author declares no competing interests.

---

**Figure S1.**
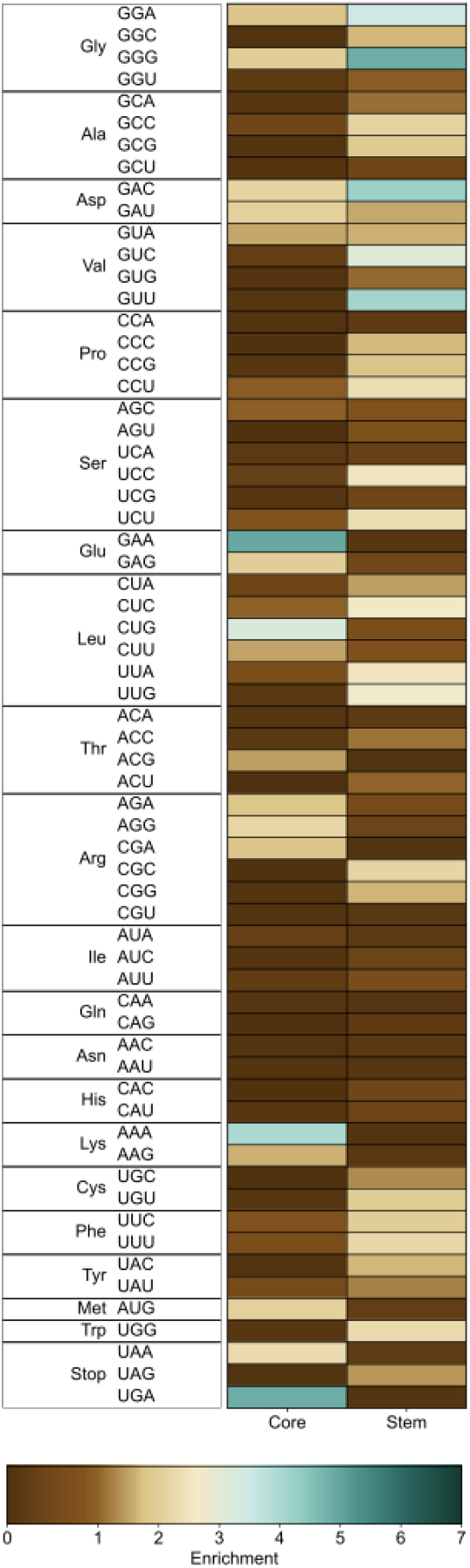
Partitioning of amino acid roles and enrichment of stop codons in catalytic cores across small, self-cleaving ribozymes (∼ 221,000).

## Notes

### Competing Interest Statement

The authors have declared no competing interest.

